# Evaluation of optically tailored fluorescent silicon quantum dots for bioimaging of the tear film

**DOI:** 10.1101/2020.12.09.411876

**Authors:** Sidra Sarwat, Fiona Jane Stapleton, Mark Duncan Perry Willcox, Peter B O’Mara, Richard D Tilley, J. Justin Gooding, Maitreyee Roy

**Affiliations:** School of Optometry and Vision Science, University of New South Wales, Sydney, Australia.; School of Chemistry and the Australian Centre for Nanomedicine, University of New South Wales, Sydney, Australia

## Abstract

This experimental study aimed to investigate the feasibility of using silicon quantum dots doped with transition metals: scandium, copper and zinc as contrast agents for eventual application for the study of the tear film in eyes. Si-QDs were synthesized and characterized by transmission electron microscopy, photoluminescence, absorbance and transient absorption measurements. The fluorescence of Si-QDs was investigated when combined with TheraTears^®^ (a balanced electrolyte formula for dry eye therapy). An optical imaging system composed of a modified slit lamp biomicroscope combined with a high-resolution Zyla sCMOS camera, SOLIS software, custom-made optical mounts and emission filters (460 nm, 510 nm and 530 nm) were used for *in vitro* imaging of Si-QDs with TheraTears^®^. The average size of Si-QDs was 2.65 nm. *In vitro* imaging of Sc-Si-QDs and Cu-Si-QDs indicated their stable and bright fluorescence with TheraTears^®^. Sc-Si-QDs were significantly brighter compared to Cu-Si-QDs and Zn-Si-QDs, and the Zn-Si-QDs showed a tendency to clump in TheraTears^®^. The fluorescence of the Si-QDs was detected down to a concentration of 0.01 µg/mL within a total volume of 5 µL. Cu-Si-QDs and Sc-Si-QDs showed brighter fluorescence than Zn-Si-QDs. However, Zn-Si-QDs and to a lesser extent, Cu-Si-QDs showed some aggregation at specific concentrations. Sc-Si-QDs are proposed as a better option for further development as an *in vivo* bioimaging agent to study the tear film dynamics.

## Introduction

Dry eye disease (DED) is one of the most common ocular conditions worldwide and commonly reported reason for seeking eye care.^1, 2^ According to the TFOS DEWS II epidemiology report, the prevalence of DED ranges from 5 to 50% in individuals over the age of 50, and it increases with age, sex and Asian ethnicity.^3^ Biochemical and biophysical changes in the tear film are associated with DED.^4^ The tear film is a dynamic fluid covering the anterior ocular surface,^5^ consisting of an outer lipid layer with an underlying muco-aqueous layer.^6^ However, how these changes impact clinical signs and tear film stability are still unknown^6^. Different models have been proposed for changes to the structure and function of the tear film in DED.^7, 8^ For instance, fluorescence microscopy with improved resolution and sensitivity has enabled labelling and visualization of proteins and lipids in live cells to understand inter- and intra-cellular dynamics.^9^ However, fluorescence microscopy has two main challenges; biological autofluorescence in the visible spectrum, and photobleaching, hence, compromising their long term fluorescence imaging.^10, 11^ In addition, the use of a relatively large volume of fluorescein (10–20 μL) applied to the total 4 μL of the non-stimulated tear film,^12^ may disrupt the structure and interfacial interactions of the tear film.^13^ To overcome the drawbacks of imaging techniques and organic dyes, quantum dots (QDs) have been used to examine the dynamics of the tear film.^13^ Commercially available indium-phosphide-gallium QDs with a zinc sulfide shell were functionalized with either hydrophilic or hydrophobic surface molecules to study tear film dynamics.^13^ One of the issues with these QDs is the cytotoxicity of indium and gallium.^14^ QDs (≤20 nm) behave as functional units comparable to the size of peptides and drugs.^15^ QDs have been used as labelling agents^16^ as they are bright, offer near-infrared region fluorescent emission and greater photostability than many organic fluorophores.^17^ The high photoluminescence quantum yield of QDs makes them good candidates for fluorescence bioimaging.^18^ Cadmium selenium/zinc sulfide QDs are the most common QDs and their emission wavelength can be tuned throughout the visible and near-infrared region of the electromagnetic spectrum,^19^ but there are concerns with the toxicity of cadmium containing QDs.^20^ The potential biocompatibility of silicon makes photoluminescent silicon QDs (Si-QDs) an ideal candidate for fluorescence imaging and may eliminate any potential toxicological problems.^21^ Studies show that silicon nanomaterials are biocompatible with human corneal epithelial cells.^22-24^ Doping of Si-QDs with transition elements enables a wide range of emission tunability and enhanced fluorescence emission intensity.^25^ Si-QDs doped with copper (Cu) have been used as near-infrared luminescent probes for the detection of heavy metals in biological systems.^26^

This study outlines the synthesis of Si-QDs doped with three different transition metals (copper, scandium, and zinc) and their characterization based on the size distribution and photoluminescence and *in vitro* imaging in TheraTears^®^. Scandium (Sc) and zinc (Zn) have been used as dopants for the first time to determine their effect on the optical characteristics of Si-QDs in comparison with copper (Cu). Zn and Sc are transition elements that share the 4^th^ period with Cu in the periodic table. Hence, Zn and Sc are expected to induce similar fluorescence in Si-QDs. Cu-Si-QDs, Sc-Si-QDs and Zn-Si-QDs were examined for the stability of their fluorescence detection limit over time by *in vitro* imaging with TheraTears^®^.

## Experimental

### Synthesis of Si-QDs

Si-QDs doped with two or four dopant atoms per particle (Cu, Sc and Zn) were synthesized by adding 0.5 g of tetraoctylammonium bromide (Sigma Aldrich, Australia) and 0.026 mmol of anhydrous salt such as ScCl_3_, CuCl_2_ or ZnCl_2_ (Sigma Aldrich, Australia) to a Schlenk tube. The Schlenk tube was then attached to a Schlenk line for the triple cycle of evacuation and purging with nitrogen for 5 minutes per cycle. 50 mL of anhydrous toluene (Sigma Aldrich, Australia) was then added, and the mixture was stirred for 24 hours. Silicon tetrachloride (Sigma Aldrich, Australia) was added to the mixture, and this was stirred for an hour. Five equivalents of lithium aluminium hydride (LiAlH_4,_ a reducing agent) (Sigma Aldrich, Australia) were added, and this was left to react for 3 hours. This procedure yielded hydride capped Sc-, Cu- or Zn-Si-QDs. Excess lithium aluminium hydride was quenched using ethanol added dropwise (until no bubbles formed).

### Surface passivation of Si-QDs

The surface of the hydride capped Si-QDs was passivated by adding anhydrous allylamine to produce hydrophilic surfaces.^27^ A quartz tube was attached to the Schlenk line and the solution degassed by triple cycles of evacuation and purging with nitrogen for 5 minutes per cycle. Hydride-capped Si-QDs were transferred via degassed syringe from the Schlenk tube to the quartz tube. Degassed anhydrous allylamine was added to the mixture, and the tube was exposed to UV light for 4 hours giving propylamine (hydrophilic) capped Si-QDs.

### Purification of Si-QDs

Purification of Si-QDs is important for *in vivo* bioimaging, where unreacted material or side products have toxic effects.^28^ After passivation, propylamine-Si-QDs were transferred to a round bottom flask, and the solvent was removed under reduced pressure. Then, Milli-Q water was added, and the mixture was dispersed by ultrasonication for 5 minutes resulting in a cloudy white solution. This mixture was filtered through a 0.45 μm filter. The resulting filtrate was concentrated to 2-3 mL under low pressure and poured into a size exclusion column containing Sephadex LH-20 beads (GE Lifesciences, Australia). The fractions were collected in an automated test tube collector and checked for luminescence using handheld UV light (365 nm). The luminescent fractions were further concentrated under reduced pressure to yield purified hydrophilic Si-QDs.

### Characterization of Si-QDs

The samples were prepared for transmission electron microscopy (OLYMPUS Life Sciences, Australia) by drop-casting the purified doped Si-QDs suspended in 0.5-1.0 mL of ethanol for hydrophilic Si-QDs on carbon-coated copper grids. TEM images were taken at an acceleration voltage of 200 kV. Photoluminescence of each type of QD was recorded on a spectrofluorophotometer (Shimadzu, Australia), using an excitation and emission slit width of 3 nm. A UV-vis spectrometer (Agilent, USA) was used to record absorbance of the Si-QDs.

### Development of an optical imaging system

A slit lamp biomicroscope (Carl Zeiss, Germany); commonly used instrument to examine the human eye, was modified with a high-resolution 5.5-megapixel Zyla sCMOS camera (Andor Technology Ltd, UK), custom-made optical mounts and emission filters (460 nm, 510 nm and 530 nm) to capture the images of fluorescence emission from Si-QDs. The Zyla sCMOS camera can split the detection path, thus allowing 100% of the reflected light to enter the camera. This camera offered full spectrograph, automatic spectral line identification and camera control along with 2D and 3D data acquisition. High-quality data export options were available as were two main (vertical and horizontal) binning variants to give binning patterns. SOLIS software (Andor Technology Ltd, UK) was used to control the camera and capture the images. Custom made optical mounts were built by the workshop staff at the Faculty of Science at the University of New South Wales. Optical mounts were designed to be comparable in size to the microscope slides used (Figure 1). Emission filters (480 nm) were placed precisely in front of the objective lens with the help of a sliding optical mount.

**Figure 1.**
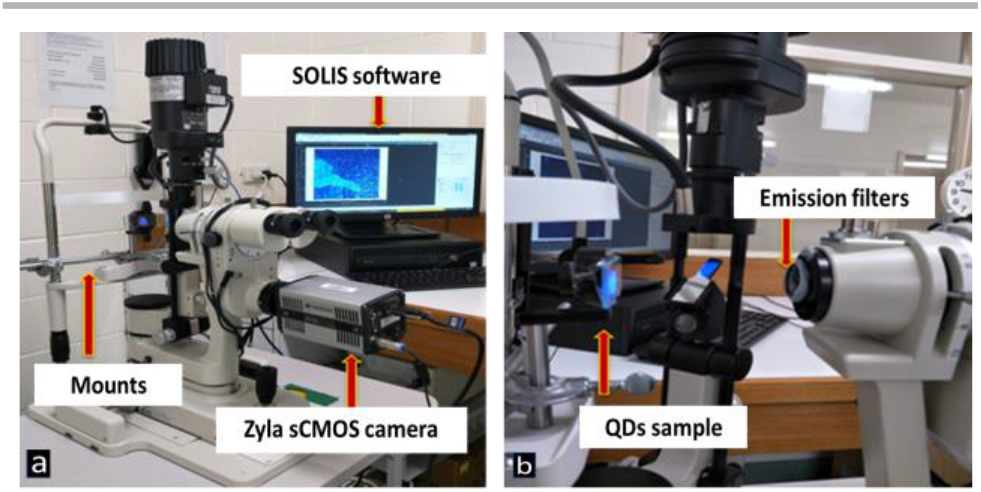
(a & b) Components of an optical imaging system.

### *In vitro* imaging of Si-QDs

Fluorescence of the Si-QDs was explored by imaging 15 different concentrations from 0.01 μg/mL to 16 μg/mL (serial dilution method) of the Cu-Si-QDs, Sc-Si-QDs and Zn-Si-QDs in TheraTears^®^ (Akorn Consumer Health, Ann Arbor, MI, USA; a balanced electrolyte formula used as lubricating eye drops) on microscope slides. Each concentration was monitored for 20 minutes, and images were captured at five different time intervals (1, 5, 10, 15 and 20 minutes). Aliquots (5 μL) of the diluted Si-QDs were added to microscope slides. The microscope slides were then sealed with clear nail polish to prevent the evaporation of the solution. The microscope slides were placed precisely at the level of eyes on the chin rest of a slit lamp with the help of the optical mounts (Figure 1). The fluorescence of Si-QDs was monitored with an optical imaging system, as shown in figure 1. Images were taken with the Zyla s sCMOS camera at a frame rate of 25 per second with the increased magnification of the slit lamp (7.5x, 16x, and 35x) every 5 minutes. A clear microscope slide and TheraTears^®^ were used as controls. Background autofluorescence from the TheraTears^®^ was subtracted from the fluorescence value of Si-QDs prior to data analysis. The excitation filter incorporated in the slit lamp was used, while external emission filters 460 nm, 510 nm and 530 nm were used to compare the fluorescence emission intensities of the Si-QDs.

### Statistical analysis

Normality of data was assessed using the Shapiro-Wilk test. Differences between groups were examined using the Kruskal Wallis test with *post-hoc* comparisons using Dunn-Bonferroni correction. Significance was determined at *p*<0.05.

## Results and discussion

### Size Distribution of Si-QDs

Si-QDs may be used as a biological marker to study the dynamics of the tear film due to their optimal fluorescence, biocompatibility, and the ability to modify their surfaces. In the current study, Si-QDs were synthesized in the solution phase^27^ to produce hydrophilic Cu-Si-QDs, Sc-Si-QDs and Zn-Si-QDs. A strong reducing agent (LiAlH_4_) yielded small-sized (∼2.65 nm) nanocrystals which are desirable for quantum yield effect for fluorescence emission.^28^ Sc-Si-QDs were 2.7±0.4 nm in size (Figure2). Hydrophilic Cu-Si-QDs were also 2.7±0.4 nm in size whilst the Zn-Si-QDs had an average size of the 2.6±0.3 nm. Their surface was modified to be hydrophilic, making them suitable for specific labelling of the aqueous layer of the tear film.^13^ The attachment of functional groups can also affect the solubility in an aqueous environment, hence compromising photoluminescence.^29^ The propylamine capped silicon nanocrystals have been found accumulated effectively in the lysosome.^30^ Therefore, amine capped Si-QDs have been synthesized and are expected to disperse in TheraTears^®^ for effective *in vitro* imaging.

**Figure 2.**
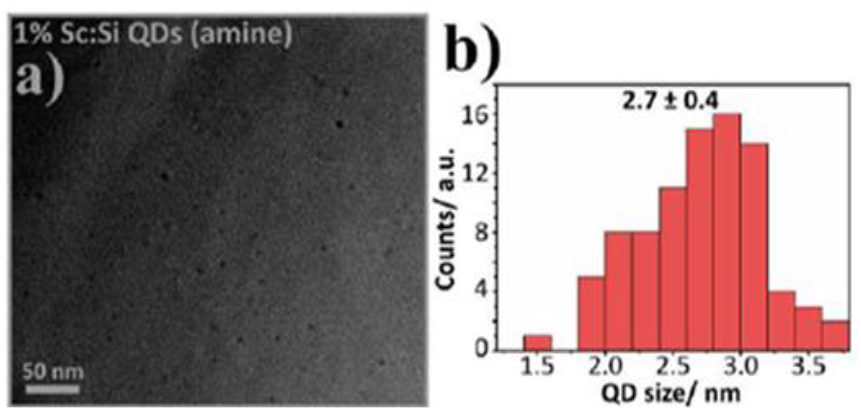
TEM images and size distribution of Sc- Si-QDs. The average size of amine and hexane capped Scandium doped Si-QDs indicates their optimal size for inducing fluorescence.

### Optical characteristics of Si-QDs

Functionalized surface-modified QDs have been used for various bioimaging studies;^15, 17, 31, 32^ however, there is only one study where QDs have been used for fluorescence imaging of the tear film of human eyes.^13^ The fluorescence emission of Cu-QDs has already been explored to image cancer cells in mice.^33^ However, the fluorescence emission of Sc-Si-QDs and Zn-Si-QDs were compared with Cu-Si-QDs for the first time in the current study by *in vitro* imaging with TheraTears^®^. The intensity of photoluminescence emission observed in Si-QDs is consistent with the previous studies.^34-36^ Figure 3 (A & B) shows the photoluminescence emission spectra of Cu-Si-QDs doped with 2 and 4 dopant copper atoms per Si-QDs using different excitation wavelengths (400-500 nm in 20 nm increments). The fluorescence emission can be enhanced by increasing the number of dopants per QDs.^25^ The dopant amount of more than two atoms per Si-QDs results in an increase in fluorescence emission intensities due to enhanced quantum yield effect.^36^ In the current study, the emission spectrum of Cu-Si-QDs with 4 dopants shows a red-shifted (shift away from UV-blue region) broadband, with high emission intensity, when using an excitation wavelength of 400 nm in contrast to 2 Cu dopants per Si-QDs. A similar effect on red-shifted broad emission wavelengths but with reduced emission intensity was observed at different excitation wavelengths. Emission results show that the optical properties such as emission wavelengths of Cu-Si-QDs may be tuned by altering the dopant concentration. Therefore, Si-QDs with 4 dopants per Si-QDs have been used for *in vitro* imaging with TheraTears^®^.

**Figure 3.**
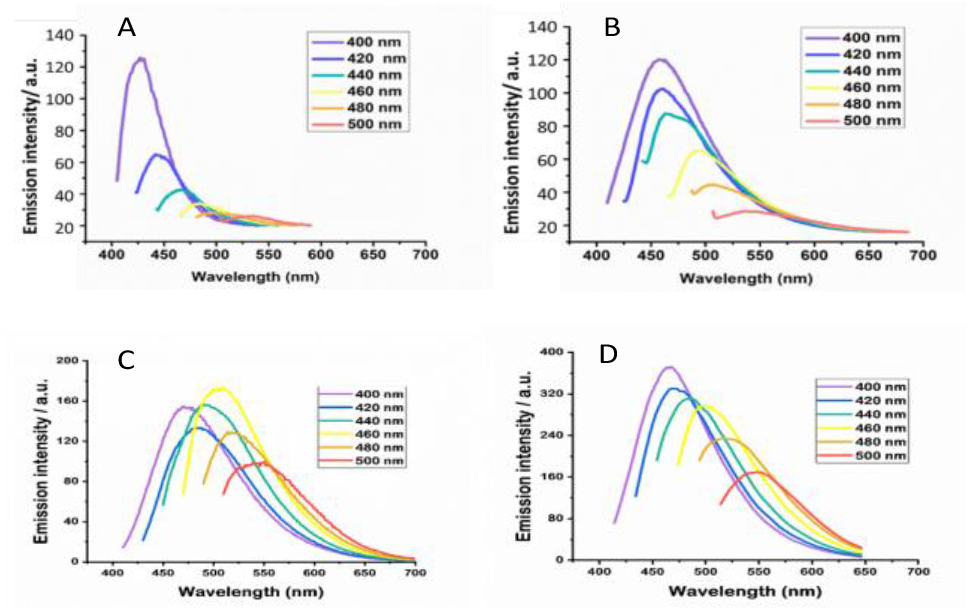
(A) Emission spectrum of Cu-Si-QDs using 2 Cu dopants per Si-QDs at different excited wavelengths (400-500 nm in 20 nm increments). (B) Emission spectrum of Cu-Si-QDs using 4 Cu dopants per Si-QDs at different excited wavelengths (400-500 nm) (c) Emission spectrum of Sc-Si-QDs at different excitation wavelengths (400-520 nm). (D)Emission spectrum of Zn-Si-QDs at different excited wavelengths (320- 600 nm).

Si-QDs showed a broad absorption range of 400-590 nm and narrow emission range (400-460 nm), which is desirable for bright fluorescence.^36^ Si-QDs are commonly excited by UV light.^29^ However, the excitation wavelength in this range causes autofluorescence and phototoxicity during bioimaging.^21^ Although a range of wavelengths that included UV was used for excitation, significant emission intensities were only seen using the visible light range, and so in practice, visible light would be preferably used for fluorescence of the QDs. The excitation range of Si-QDs was 400-590 nm, which lies in the visible range of the electromagnetic spectrum, and allowed the use of a visible light source slit lamp biomicroscope for *in vitro* imaging of TheraTears^®^ as previously used for QDs bioimaging.^13^ Furthermore, based on the optical characteristics, hydrophilic Si-QDs were found suitable for bright and stable fluorescence emission. However, hydrophobic Si-QDs need to be investigated for their optimal photoluminescence emission for the lipid layer of the tear film. Figure 3 (C) shows the photoluminescence emission spectra of hydrophilic Sc-Si-QDs using a series of excitation wavelengths from 360 nm to 520 nm in 20 nm increments. As can be seen, the photoluminescence emission spectra were dependent on excitation wavelengths and shifted toward the longer wavelengths with narrow emission bandwidth and reduced emission intensities. The maximum photoluminescence emission peak of Sc-Si-QDs was at an excitation wavelength of 460 nm, and the minimum photoluminescence emission intensity occurred at 520 nm excitation wavelength. The higher excitation wavelength has shifted fluorescence emission peak towards longer wavelength of the spectrum, but with reduced fluorescence intensity. The results show the optimal peak fluorescence emission at the 450 nm when excited at 400 nm. Zn-Si-QDs showed similar photoluminescence and absorption pattern to Cu-Si-QDs and Sc-Si-QDs. Figure 3 (D) shows the photoluminescence intensities emission spectra of Zn-Si-QDs using different excitation wavelengths from 320 nm to 600 nm in 20 nm increments. As Cu-Si-QDs and Sc-Si-QDs, the photoluminescence emissions are dependent on the excitation wavelengths. Hence the photoluminescence emission spectrum shifts toward the red and near-infrared progressively with the longer excitation wavelengths.

### *In vitro* fluorescence imaging of Si-QDs

*In vitro* fluorescence imaging of Si-QDs in TheraTears^®^ showed significant differences in fluorescence intensity between the three types of doped Si-QDs for all concentrations at given time points. *Post-hoc* comparisons showed that Sc-Si-QDs showed significantly brighter fluorescence emission than Zn-Si-QDs at concentrations such as 0.01 μg/mL and 1 μg/mL at the given time points. However, no difference in fluorescence emission was observed between Sc-Si-QDs and Cu-Si-QDs or Cu-Si-QDs and Zn-Si-QDs at concentrations of 16 μg/mL and below. At higher concentration of 16 μg/mL, Cu-Si-QDs and Zn-Si-QDs were significantly different in fluorescence emission for given time points; however, no difference in fluorescence intensity was observed between Sc-Si-QDs and Zn-Si-QDs. Again, no difference was observed between Sc-Si-QDs and Cu-Si-QDs at the higher concentrations. One surprising result was the difference between Cu-Si-QDs and Zn-Si-QDs at 1 μg/mL and between Zn-Si-QDs and Sc-Si-QDs at 16 μg/mL after 15 mins. Overall, Sc-Si-QDs gave higher fluorescence emission than Zn-Si-QDs at lower concentrations, and Cu-Si-QDs gave higher emissions than Zn-Si-QDs at higher concentrations. The fluorescence emission of Cu-Si-QD and Sc-Si-QDs were not significantly different at any concentration.

Similarly, figure 4 shows that the control (TheraTears^®^) without Si-QDs gave no fluorescence while Si-QDs had varying fluorescence intensities for given concentrations at different time points. Cu-Si-QDs and Sc-Si-QDs provided bright fluorescence signals, while Zn-Si-QDs showed dispersed fluorescence emission at 16 μg/mL. Fluorescence of three types of Si-QDs was detectable at all concentrations tested. The fluorescence intensity of Sc-Si-QDs was less bright than that of Cu-Si-QDs at 16 μg/mL and above (Figure4); however, there was no statistically significant difference in signal intensity (*p* 0.0001). Zn-Si-QDs and Cu-Si-QDs were seen to aggregate during *in vitro* imaging (arrows Figure 4). Although aggregation has been reduced with decreasing concentration, it was still observed at 0.01 μg/mL of Cu-Si-QDs and Zn-Si-QDs. Zn-Si-QDs appeared to aggregate more than Cu-Si-QDs, and Sc-Si-QDs showed no aggregation at any concentration (figure 6). Zn-Si-QDs aggregated at 0.01 μg/mL, 1 μg/mL and 16 μg/mL, and this has been reported previously as an issue with QDs.^37^ This may be due to their functional surface groups (propylamine), which might have shown attraction towards the aqueous components in TheraTears^®^. The aggregation of QDs can be addressed by dispersion in a suitable surfactant^38^ as it provides stability and uniform dispersion.^37^ However, surfactants may not be useful for *in vivo* imaging as they disrupt the tear film.^39^ Encapsulation and surface modification are the most common methods used to increase stability and inertness of QDs.^40^ Therefore, the modification of Si-QDs with functional groups such as phospholipids may enhance their dispersion in tear film lipid layer to avoid the aggregation during bioimaging. Cu-Si-QDs, Sc-Si-QDs and Zn-Si-QDs gave appreciable fluorescence in TheraTears^®^ even at concentrations as low as 0.01 μg/mL. Cu-Si-QDs and Sc-Si-QDs showed brighter fluorescence as a function of time compared to Zn-Si-QDs.

**Figure 4.**
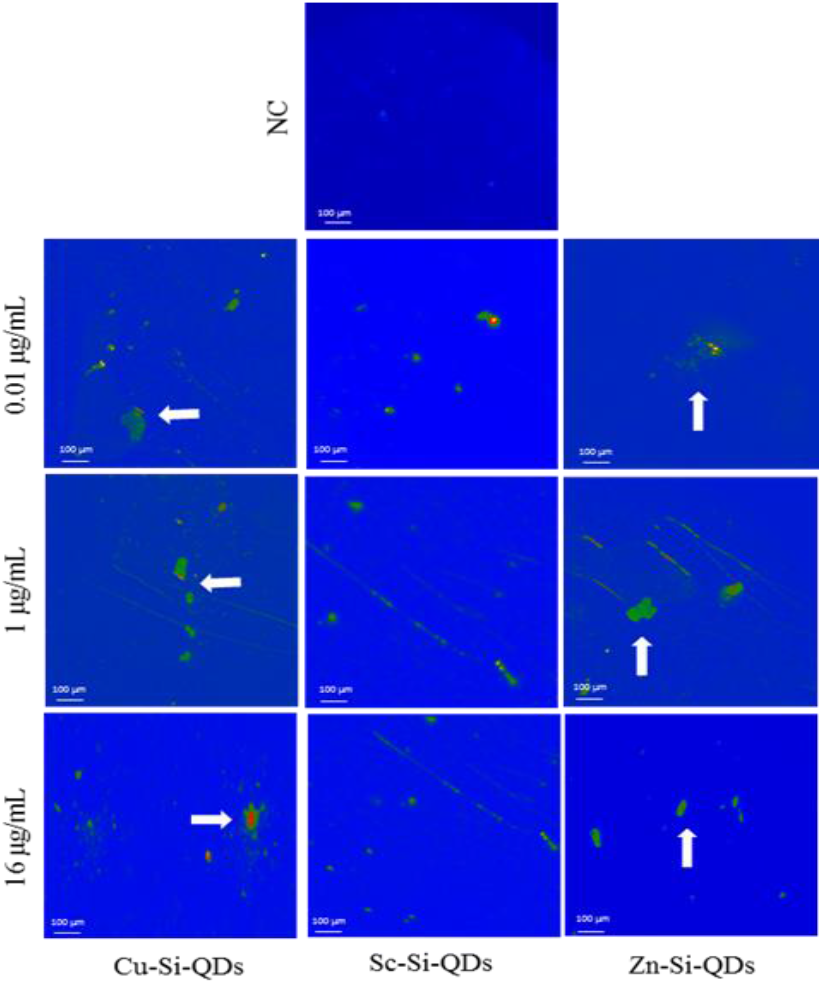
Fluorescence emission of Cu-Si-QDs, Sc-Si-QDs and Zn-Si-QDs at different concentrations. NC= negative control (TheraTears alone) and arrowheads show the aggregation of Si-QDs.

Figure 5 shows time series of fluorescence emission for the Cu-Si-QDs, Sc-Si-QDs and Zn-Si-QDs at 4 different concentrations (0.01, 1, 8 and 16 μg/mL), that have been shown to be non-toxic (data not shown here), at 5 different time points (1, 5, 10, 15 and 20 minutes). This figure shows no significant difference in fluorescence emission among Sc-Si-QDs and Cu-Si-QDs at different time points (*p*<0.0001). Sc-Si-QDs showed the highest fluorescence emission, even at the lowest concentration of 0.01 μg/mL. Zn-Si-QDs emit less intense fluorescence emission compared to both Sc-Si-QDs and Cu-Si-QDs at all time points, but fluorescence emission was almost equivalent to Sc-Si-QDs and Cu-Si-QDs after 20 minutes. However, Zn-Si-QDs showed similar fluorescence emission intensity at a minimum concentration of 0.01 μg/mL. Overall, the fluorescence emission of Si-QDs remained stable for 20 minutes, hence making Si-QDs suitable for bioimaging of tear film and studying its dynamics for longer period.

**Figure 5.**
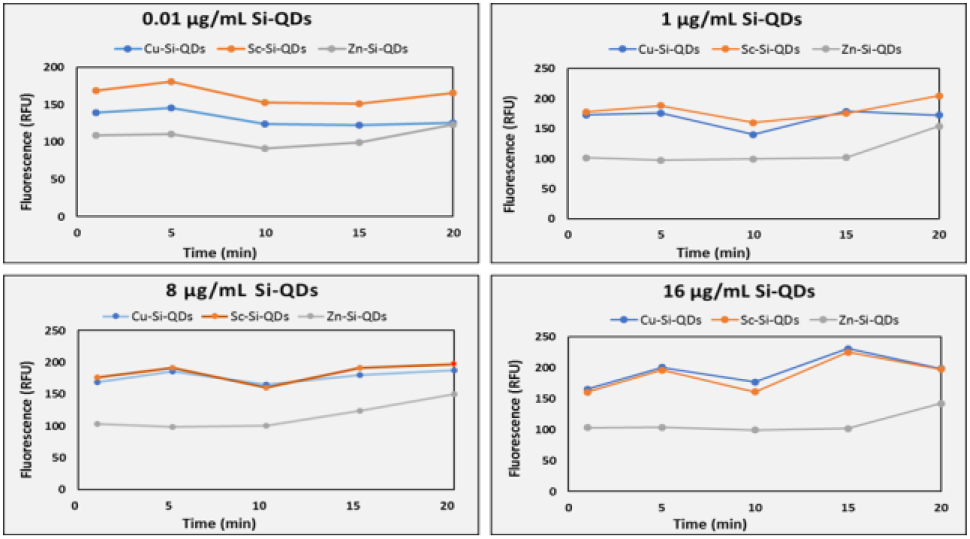
Time series of fluorescence emission of Cu-Si-QDs, Sc-Si-QDs and Zn-Si-QDs for 4 different concentrations (0.01, 1, 8 and 16 μg/mL) at 5 different time points. (1, 5, 10, 15 and 20 minutes). NS: Non-significant. RFU: Relative fluorescence unit.

## Conclusions

Sc-Si-QDs, Cu-Si-QDs and Zn-Si-QDs have been successfully synthesized and investigated for their optimal photoluminescence characteristics. Si-QDs with the size range of 2-4 nm are optimal for fluorescence emission; therefore, small-sized Si-QDs approximately 2.7 nm with 4 dopants transition elements will be effective for fluorescence imaging of the tear film. The fluorescence detection limit demonstrates the possible use of Cu-Si-QDs and Sc-Si-QDs for *in vivo* imaging of tear film, as Zn-Si-QDs have reduced fluorescence intensity comparatively and tend to aggregate more. However, Cu-Si-QDs also showed some aggregation of particles in TheraTears^®^. Therefore, Sc-Si-QDs appeared to be a better option for future *in vivo* bioimaging of tear film, but the real challenge is to deliver the hydrophobic Si-QDs to the lipid layer of the tear film without using a toxic organic solvent like hexane, which needs further research.

## Author Contributions

These authors contributed equally to this work.

## Acknowledgements

This project was funded by a University of New South Wales Faculty Research Grant and Faculty of Science Interdisciplinary Grant.

